# Inference of bacterial small RNA regulatory networks and integration with transcription factor driven regulatory networks

**DOI:** 10.1101/657478

**Authors:** Mario L. Arrieta-Ortiz, Christoph Hafemeister, Bentley Shuster, Nitin S. Baliga, Richard Bonneau, Patrick Eichenberger

## Abstract

Small non-coding RNAs (sRNAs) are key regulators of bacterial gene expression. Through complementary base pairing, sRNAs affect messenger RNA stability and translation efficiency. Here, we describe a network inference approach designed to identify sRNA-mediated regulation of transcript levels. We use existing transcriptional datasets and prior knowledge to infer sRNA regulons using our network inference tool, the *Inferelator*. This approach produces genome-wide gene regulatory networks that include contributions by both transcription factors and sRNAs. We show the benefits of estimating and incorporating sRNA activities into network inference pipelines. We comprehensively assess the accuracy of inferred sRNA regulons using available experimental data. We uncover 30 novel experimentally supported sRNA-mRNA interactions in *Escherichia coli*, outperforming previous network-based efforts. Our findings expand the role of sRNAs in the regulation of chemotaxis, oxidation-reduction processes, galactose intake, and generation of pyruvate. Additionally, our pipeline complements sequence-based sRNA-mRNA interaction prediction methods by adding a data-driven filtering step. Finally, we show the general applicability of our approach by identifying novel, experimentally supported, sRNA-mRNA interactions in *Pseudomonas aeruginosa* and *Bacillus subtilis*. Overall, our strategy generates novel insights into the functional implications of sRNA regulation in multiple bacterial species.

**IMPORTANCE:** Individual bacterial genomes can have dozens of small non-coding RNAs with largely unexplored regulatory functions. Although bacterial sRNAs influence a wide range of biological processes, including antibiotic resistance and pathogenicity, our current understanding of sRNA-mediated regulation is far from complete. Most of the available information is restricted to a few well-studied bacterial species; and even in those species, only partial sets of sRNA targets have been characterized in detail. To close this information gap, we developed a computational strategy that takes advantage of available transcriptional data and knowledge about validated and putative sRNA-mRNA interactions. Our approach facilitates the identification of experimentally supported novel interactions while filtering out false positives. Due to its data-driven nature, our method emerges as an ideal strategy to identify biologically relevant interactions among lists of candidate sRNA-target pairs predicted *in silico* from sequence analysis or derived from sRNA-mRNA binding experiments.

## INTRODUCTION

Although bacterial gene regulation has been primarily investigated at the transcription level, recent studies have confirmed the importance of small non-coding RNAs (sRNAs) as post-transcriptional regulators (1–5). Via complementary base pairing to their targets, bacterial sRNAs regulate transcript processing, stability and translation into proteins (3–5). sRNA binding promotes conformational changes in mRNA secondary structure thus modulating recognition by molecular complexes such as ribosomes and ribonucleases (3). Chromosome-encoded sRNAs can be classified as either trans-encoded (when they regulate genes regardless of their chromosomal location) or cis-encoded (when they solely regulate the expression of adjacent genes) (3, 6). Here, we focus on trans-encoded sRNAs affecting mRNA stability. Importantly, the list of sRNA-controlled cellular functions is broad (ranging from metabolism to virulence) and is continuously expanding with the analysis of new microbial species (5, 7). Because transcription factors (TFs) and sRNAs can share targets or even regulate each other (7), a comprehensive characterization of any bacterial gene regulatory network must incorporate both types of regulators (as has been explored for regulatory networks in eukaryotes) (8).

The role of trans-encoded sRNAs has been mainly investigated in Gram-negative bacteria (5). *Escherichia coli* is currently the bacterial species with the highest number of experimentally supported sRNA-mRNA interactions (102 known interactions according to Pain *et al*., 2015) (9). This set contains 22 sRNAs with at least one experimentally supported target (9); however, this is only a fraction of the array of potential regulatory RNAs encoded in the *E.coli* genome (10, 11). The number of characterized sRNA targets is unevenly distributed as only eight out of 22 sRNA regulons contain five or more members. Accurate and comprehensive detection of sRNA-mRNA interactions is challenging. The outcome of transcriptomics and proteomics experiments is highly dependent on the proper selection of conditions in which sRNAs are regulatory active (12), further complicating experimental designs. Moreover, computational methods (based on genome sequence and hybridization energy) predicting sRNA-mRNA interactions are fast and inexpensive but have a high false positive rate and may fail to recall known targets (5, 9).

Network inference methods have been implemented to study sRNA-mediated regulation. Modi and collaborators used Context Likelihood of Relatedness (CLR) on transcriptional profiles of sRNAs and genes to infer an *E. coli* sRNA regulatory network (13, 14). Modi et al. correctly predicted *lrp*, encoding a global transcriptional regulator, as a target of the GcvB sRNA. A second study exploited gene co-regulation to infer another *E. coli* sRNA network (15). In both studies, the recall of known sRNA-mRNA interactions was limited and the accuracy of novel predictions was not systematically evaluated (15).

We hypothesize that, contrary to what was assumed in previous sRNA network inference strategies, sRNA levels might not be an adequate proxy for their regulatory activity in large transcriptomic datasets. In multiple species, RNA chaperones (such as Hfq in *E. coli*) promotes the interaction between sRNAs and their target mRNAs (4, 5, 16). Moreover, ribonucleases may be required to activate sRNAs by processing (e.g. RoxS, a *Bacillus subtilis* sRNA, only interacts with the *sucCD* mRNA after it has been truncated by RNase Y) (17). Furthermore, the regulatory contribution of a sRNA becomes negligible when the concentration of its targets significantly exceeds its own (18, 19). In this work, we address the complexity of sRNA-mediated regulation by estimating sRNA regulatory activities from transcriptional profiles of their known and candidate targets. We then use the estimated sRNA activities as input to our network inference tool to generate models of gene regulation for four bacterial species. We show, with substantial experimental support from independent studies, that our pipeline outperforms previous network-oriented efforts, detects novel sRNA-mRNA interactions, and complements RNA-RNA interaction prediction methods by discriminating between true and false targets. This work illustrates how our computational strategy can help researchers selecting candidate interactions for experimental validation while focusing on the most likely sRNA targets.

## RESULTS AND DISCUSSION

We inferred bacterial sRNA regulons from transcriptomics data using either the Inferelator (20), our network inference tool, or CLR, an alternative algorithm (13, 21). Because our approach mines transcriptomics data, it is designed to identify sRNA-mRNA interactions that change mRNA stability (those that only modify translational efficiency would likely be overlooked). A set of experimentally supported sRNA-mRNA interactions (also referred to as sRNAs priors) was used for estimating sRNA regulatory activities (*see below*). We used *E. coli* data to benchmark our pipeline and restricted our analysis to eight sRNAs with experimentally supported targets (**Table 1**). We repeated this strategy with *B. subtilis*, *Staphylococcus aureus* and *Pseudomonas aeruginosa*. sRNA priors used for estimating sRNA activities are listed in **Table S1**. We relied on publicly available experimental data for assessing the accuracy of the inferred sRNA regulons.

**Table 1.** Escherichia coli sRNAs analyzed in this study

| sRNA | Biological process | Prior Targets | Differentially expressed genes<sup>\$</sup> | References |
| --- | --- | --- | --- | --- |
| CyaR | Sugar metabolism | *luxS, nadE, ompX, yqaE, yobF, ptsI* | 28 | (63) |
| FnrS | Anaerobic respiration | *cydD, folE, folX, gpmA, maeA, marA, metE, sodA, sodB, yobA* | 59 | (61, 62) |
| GcvB | Amino acid metabolism & transport | *argT, csgD, cycA, gdhA, livJ, lrp, phoP, sstT, yifK* | 88 | (64) |
| MicA | Stress response | *ecnB, fimB, lamB, lpxT, ompA, ompW, tsx, ycfS, yfeK* | 16 | (65) |
| OmrA/OmrB<sup>^</sup> | Stress response (membrane) | *cirA, csgD, fecA, fepA, opmR, opmT* | 48 | (66, 67) |
| RybB | Stress response | *fadL, fiu, lamB, nmpC, ompA, ompC, ompF, ompW, rluD, tsx* | 22 | (65) |
| RyhB | Iron metabolism | *acnA, cysE, dmsA, erpA, fumA, fumB, msrB, nagZ, sodB, uof, ykgJ, ynfF* | 87* | (24, 31) |
| Spf | Sugar metabolism & transport | *ascF, fucl, galk, glpF, gltA, maeA, nanC, paaK, puuE, srlA, sthA, xylF* | 42<sup>#</sup> | (37) |

<sup>\$</sup>Based on available profiling data
\*Includes ribosome profiling data
<sup>#</sup>Re-analyzed using Cyber-T (59)
<sup>^</sup>OmrA and OmrB were merged in a single regulator (OmrA)

### sRNA transcript level is not a good proxy for regulatory activity in a network inference context

The transcriptional profiles of sRNAs have commonly been used as proxies for their regulatory activities (14, 15). However, we suspected that a sRNA transcriptional profile would not typically match its regulatory activity due to the contributions of factors (such as RNA chaperones, ribonucleases, RNA sponges, target mRNA concentration) that influence the outcome of sRNA-mediated regulation. An analogous observation has been made for TFs, where TF activity can be modulated by post-translational modifications such as phosphorylation or the presence of co-factors (22, 23). To examine the relation between the transcription level of a sRNA and its regulatory activity in *E. coli*, we plotted the transcriptional profile of several sRNAs against the average transcription profile of their experimentally supported targets (**Fig. 1A-B & Fig. S1 A-E**).

**Figure 1.**
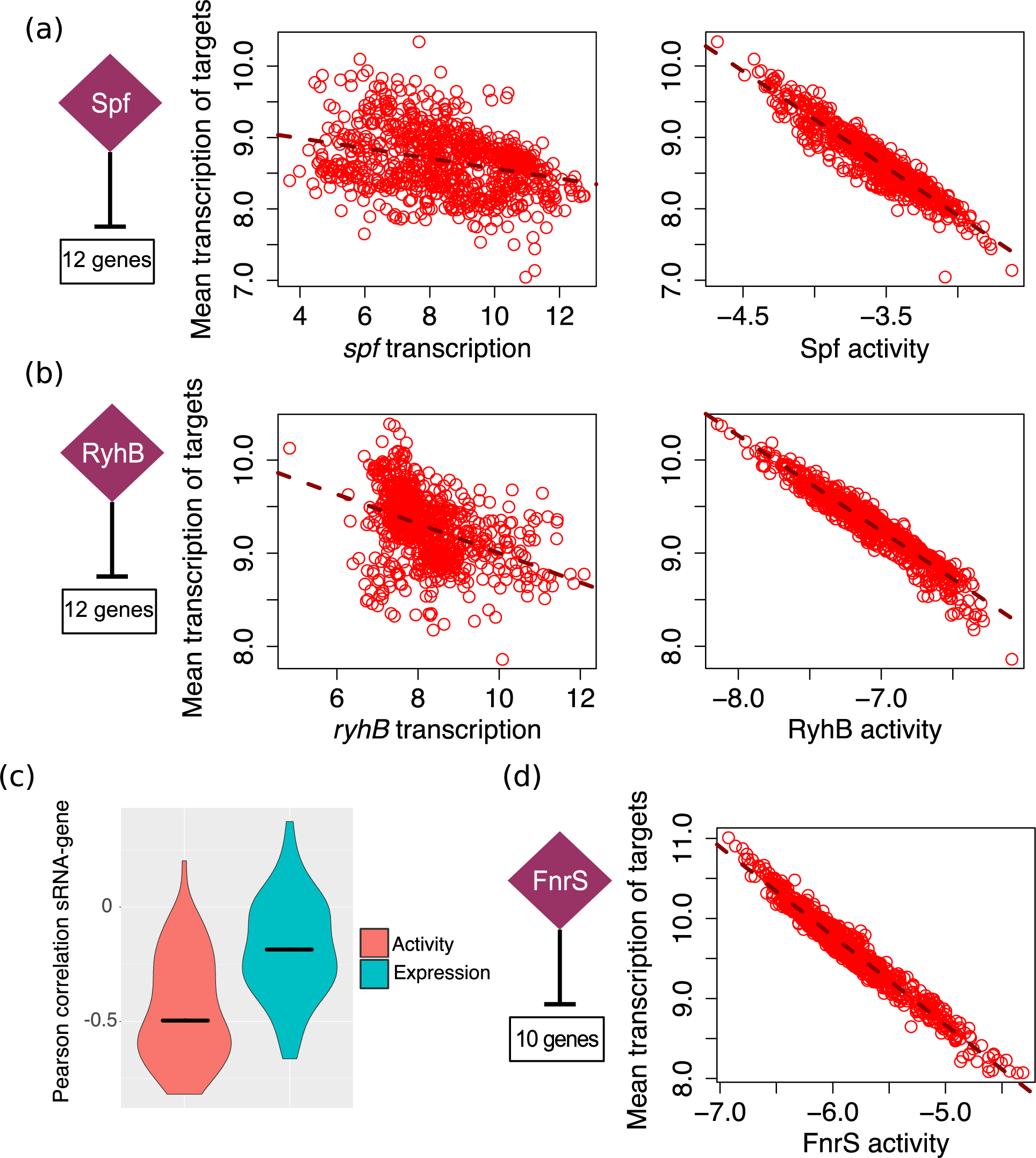
The transcriptional profile of a sRNA is a sub-optimal proxy for its regulatory activity. The motivation for estimating sRNA activities is illustrated for three *E. coli* sRNAs. sRNA activities were estimated for each experimental condition. Each dot represents one microarray experiment. The number of known targets used to estimate sRNA activities and to compute the mean transcription of the analyzed regulons (in each condition) is indicated. A) Spf controls the uptake and metabolism of alternative sugars (37). A stronger relation is observed between the estimated Spf activity and the mean transcription profile of its dependent genes (right panel) than between the transcription profile of *spf* and its targets (left panel). B) RyhB is involved in iron metabolism and represses expression of iron-consuming genes as part of the iron sparing response under iron poor conditions (31). Similarly, the relation between estimated RyhB activity and the mean transcription profile of its targets is stronger than the relation between the transcription profile of *ryhB* and its targets. C) Violin plots show the distribution of Pearson correlation values between sRNAs and the transcriptional profile of their priors when either estimated sRNA activities or sRNA transcriptional profiles are used for computation. Black lines indicate median correlation values (−0.5 and −0.18 for sRNA activity and sRNA transcriptional profiles, respectively). The difference between both sets of correlation values is statistically significant (T-test p-value = 9.3×10^−10^). D) FnrS is involved in respiration (61, 62). Probes for the *fnrS* gene did not need to be present in the *E. coli* transcriptomic dataset in order to be included as potential regulator in our pipeline. FnrS activities were estimated from the transcriptional profile of 10 FnrS-dependent genes present in the dataset.

In agreement with our expectations, sRNA transcript levels exhibited only a weak linear relation with their targets (left panels). This observation holds true for other species (**Fig. S1 F-G**). For instance, we observed similar patterns for two regulators of the iron-sparing response, FsrA in *B. subtilis* and S596 in *S. aureus*, functional analogs of *E. coli* RyhB (24–26). These findings support the notion that transcript levels are often a sub-optimal proxy for sRNA regulatory activity in the context of network inference.

### Estimating sRNA regulatory activity

To estimate sRNA regulatory activity (SRA), we used the transcription profiles of their experimentally supported targets. Conceptually, this is analogous to relying on a reporter gene to measure the activity of a given sRNA, with the distinction that every presumed target of the sRNA is considered in the estimation (27). We have successfully used a similar approach to estimate the activities of TFs and thereby expanded the transcriptional network model of *B. subtilis* (27). We checked the relation between estimated SRAs and the transcription profile of their priors (**Fig. 1A-B & Fig. S1 A-E**; right panels). We observed, as expected based on our previous work (27), a stronger linear relationship between genes and their known sRNA regulators than with raw sRNA transcript levels. We noted the same trend for functionally related sRNAs in *B. subtilis* and *S. aureus* (**Fig. S1 E-G**). Additionally, significantly stronger anti-correlation (expected due to the repressive nature of sRNA-mRNA interactions used as priors) were found between sRNAs activities and their targets compared to correlations between corresponding sRNA transcript levels and their targets (**Fig. 1C**).

The better correlation between sRNA activity and transcriptional profile of sRNA targets led us to incorporate SRAs into our inference pipeline in the same manner we did for TFs. Importantly, estimated SRAs can be used for network inference even when the transcriptomic dataset does not contain information about sRNAs of interest, as frequently observed for microarrays-collected datasets. One example is shown in **Fig. 1D**. Despite the absence of FnrS in the transcriptomic dataset, its activity was estimated using ten priors. In our workflow, the only requirement for including a sRNA as potential regulator is a set of experimentally supported or candidate targets, whose transcriptional profiles are available in the analyzed transcriptomics dataset.

### General strategy

Our network inference pipeline is illustrated in **Fig. 2**. First, we used a transcriptomics dataset (from the Many Microbe Microarrays database (28) or any equivalent repository) and a set of experimentally supported TF-gene and sRNA-mRNA interactions (from RegulonDB (29), RegPrecise (30), or equivalent), referred to here as the prior network, to estimate the regulatory activities of TFs (TFAs) and sRNAs (SRAs). Next, we used the estimated activities (TFAs and SRAs), the transcriptomics dataset, and the prior network to simultaneously infer the TF-controlled network and the sRNA-controlled network using Bayesian regression with the Inferelator (*see methods*)(20, 27). Interactions not included in the prior network were considered novel. Inclusion of a prior transcriptional network, which is much larger than the prior sRNA network, allowed us to define thresholds (calibrated using desired precision values) for selecting the interactions that should be kept in the final networks. Inclusion of TFs also prevented model over-fitting due to an incomplete set of regulators and interactions to explore. Additionally, the simultaneous inference of the transcriptional and post-transcriptional networks enabled us to study connections between the two regulatory layers.

**Figure 2.**
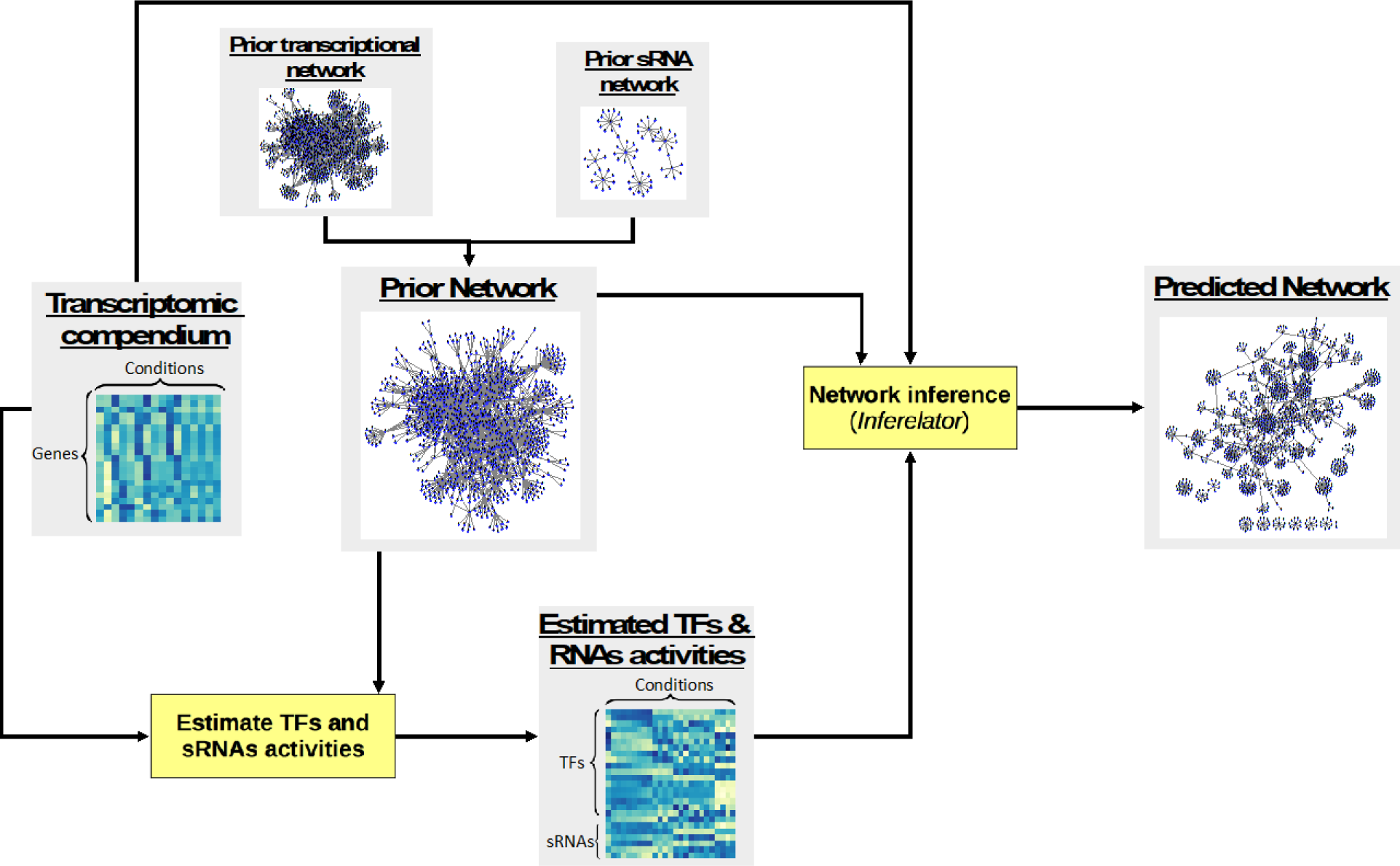
General strategy. A transcriptomic dataset and a prior network (built from experimentally supported TF-gene and supported or candidate sRNA-mRNA interactions) are used for estimating the regulatory activities of TFs (TFAs) and sRNAs (SRAs) using a network component analysis approach (27, 57). Next, estimated TFAs and SRAs, transcriptomic data and prior network are used as input for the *Inferelator* to infer a regulatory network composed of a transcriptional layer (TF-based) and a post-transcriptional layer (sRNA-based).

### Our strategy improves performance, recovers known interactions and predicts novel sRNA-mRNA interactions

We compared the performance of the Inferelator (using a *B*ayesian *B*est *S*ubset *R*egression-BSSR) and mixed-CLR, with and without incorporating sRNA activities (SRA). For each method, the number of predicted mRNA targets per sRNA versus the number of predicted targets with experimental support is shown in **Fig. 3A**. Importantly, genes used as priors for sRNA activity estimation were removed from the set of predicted targets because they tend to occupy high positions in the predictions ranking. FnrS was not considered in this analysis because its transcriptional profile was missing from the transcriptomic dataset. Thus, it cannot be included as a regulator in methods that use transcriptional profiles as proxy for activity. We deemed a predicted target to be experimentally supported if it was differentially expressed in transcriptional profiling experiments overexpressing or deleting its putative sRNA regulator (according to the criteria established in the corresponding publication, except for Spf, *see methods*). Additionally, a sRNA-mRNA interaction was considered experimentally supported when the predicted target was part of an operon that contains differentially expressed genes. For RyhB, available ribosome profiling data was also considered in evaluating experimental support (31). The sets of candidate sRNA targets identified with transcriptomic experiments contain genes whose expression is (directly or indirectly) affected by the sRNA of interest.

**Figure 3.**
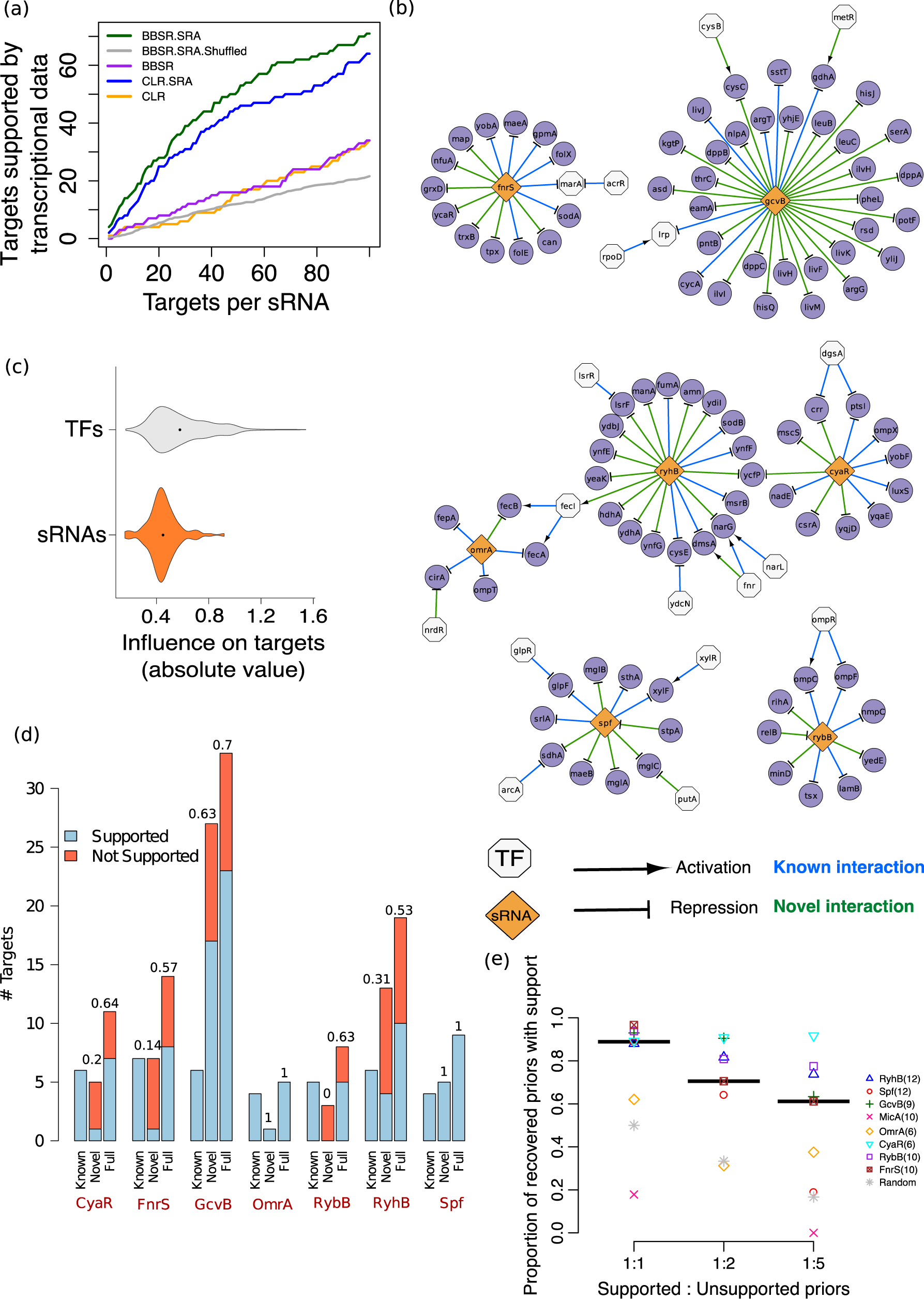
Performance of the Inferelator and alternative computational methods for expanding sRNA networks. A) Performance of the Inferelator (BBSR) and mixed-CLR, an alternative method, with (indicated by the SRA suffix) and without incorporation of sRNA activities. Genes predicted as targets but not used for sRNA activity estimation were considered to be experimentally supported if they were differentially expressed in transcriptional profiling experiments (deletion or over-expression of CyaR, GcvB, MicA, OmrA, Spf, RybB and RyhB) or when they were part of an operon containing differentially expressed genes. For each sRNA, targets were ranked based on confidence score (in the case of the Inferelator) or mutual information-based score (in the mixed-CLR runs). To estimate the basal performance level of the Inferelator, the average of ten runs with shuffled sRNA priors was also computed (grey line). B) The inferred sRNA regulatory network of *E. coli*. To allow comparison between transcriptional and post-transcriptional networks, overlap between both networks is displayed. C) Violin plots showing distribution of absolute values of Bayesian regression coefficients (which indicate magnitude and direction) associated with TF-gene and sRNA-mRNA interactions. Black dots indicate the median. D) The inferred sRNA regulons are experimentally supported. Experimental support rate for novel predictions (not in the prior network) and full inferred regulons (recovered priors and novel predictions) of the Inferelator.SRA run described in panel A are shown on top of each bar. E) The Inferelator can identify experimentally supported targets among noisy priors. Experimental support rates for recovered priors are plotted for different levels of noise in the priors. Each dot is the mean value of ten Inferelator runs (each run with a different set of false priors). Each colored symbol corresponds to one of eight sRNAs. Black lines indicate the median proportion for all eight sRNAs. Gray star indicates the expected proportion if priors included in the predicted networks were randomly selected. Number of true sRNA targets is shown in parentheses.

Therefore, the rank of differentially expressed genes in the list of predicted sRNA targets informs about the performance of our strategy (10). We ranked sRNA-mRNA interactions based on the confidence score computed by the Inferelator (*see methods*). When we analyzed the top 20 predictions per sRNA (for the seven *E. coli* sRNAs that were considered), we observed that among the 140 predictions made by the Inferelator with sRNA activities (Inferelator.SRA), 28 were experimentally supported (25 for mixed-CLR). By contrast, the Inferelator without sRNA activities only predicted eight experimentally supported targets (four for mixed-CLR). Inferelator.SRA performed best for Spf (ten supported targets in the top 20 predictions) and GcvB (nine supported targets). There was at least one supported target for all sRNAs except RybB and MicA. In general, we observed that incorporation of sRNA activities consistently improved the detection power of both network inference tools [in **Fig. 3A**: green and blue lines (with SRAs) vs. purple and orange lines (without SRAs)].

The inferred *E. coli* sRNA network from the Inferelator run with sRNA activities (BBSR.SRA) described above is shown in **Fig. 3B**. Limited overlap was observed between the inferred sRNA network and the TF network. Only 19% of sRNA-regulated genes were predicted as targets of one or more TFs. Despite 41% of genes having two or more regulators in the prior network, expression of most genes was explained as the function of a single regulator’s activity (either a TF or a sRNA). We found multiple cases in which the regulatory influence of a sRNA surpassed the estimated influence of several TFs targeting the same gene. For example, according to the prior network, *marA* is regulated by five TFs (AcrR, CpxR, Fis, Rob, SoxS) and one sRNA (FnrS). Only the interactions between *marA* and AcrR and FnrS were recalled into the final model. For genes predicted to be regulated by both TFs and sRNAs in the inferred network (*sdhA*, *ompC, cysC*, among other targets), TFs were commonly the most influential regulator. In fact, we observed that on average the influence of sRNAs on expression of their targets is subtler than the one exerted by TFs (**Fig. 3C**). This finding is in agreement with the view of sRNAs as fine tuners of gene expression (3). When we inferred an alternative model (with an Inferelator run in which sRNAs were not considered as potential regulators), 90% of the genes exclusively regulated by sRNAs in our original network (**Fig. 3B**) lacked regulatory hypotheses (data not shown). This finding underscores the importance of sRNAs for fine tuning of gene expression, and it demonstrates that inclusion of sRNAs as regulators expands the models of gene regulation in bacteria.

The accuracy of inferred sRNA regulons was assessed using experimental data from previously published studies (including transcriptional profiling, ribosome profiling, and sRNA-mRNA binding data). Experimental support for novel targets and entire sRNA regulons inferred with our strategy is shown in **Fig. 3D**. Thirty-eight sRNA-mRNA interactions from the prior network were included in the final model (for a total recall of 0.51). The average recall per sRNA regulon was 0.55 and the highest recall (1.0) was obtained for CyaR. In addition to the recovered priors, the inferred sRNA network contained 61 novel interactions. 29 out of these 61 novel predictions (0.48) were experimentally supported. Per regulon, the average experimental support for novel predictions was 0.47, which increased to 0.71 when considering both novel predictions and recovered priors. The limited increase in size for some sRNA regulons is consistent with previous observations that regulators with the lowest number of priors tend to have the lowest number of novel predictions because regulator’s activity cannot be estimated with precision (27). Failure to recall MicA targets is likely a consequence of the weak correlation between estimated MicA activity and the transcription profiles of its known targets (**Fig. S1C**). In future applications, the detection power of our pipeline will be improved by expanding the set of priors (for example, by including every gene differentially expressed in transcriptional profiling experiments). In the above analysis, we intentionally left out some of the potential sRNA targets to estimate the accuracy of our pipeline. In conclusion, integration of estimated sRNA activities in the network inference procedure greatly improves the ability to detect additional experimentally supported sRNA-mRNA interactions.

### Robustness to incorrect prior information

We originally tested our approach with a set of priors that only included experimentally supported sRNA-mRNA interactions. However, in a more realistic scenario, researchers may compile priors from heterogenous sources, and a mix of true and false interactions is expected. Previously, we showed that the Inferelator is robust to noisy priors (up to 1:10 ratio of true: false priors) (27). To confirm this result in the context of sRNAs, we assessed the robustness of our pipeline in terms of the experimental support of priors included in the final models. We added different amounts of false interactions to the sRNA priors and ran the pipeline with those noisy priors. We found that our method efficiently distinguishes true from false interactions (**Fig. 3E**). Specifically, we determined how many priors recovered as putative targets were experimentally supported. Although the total number of recovered priors is lower than in the original run without false priors (**Fig. S2**), the proportion of recovered priors with experimental support still exceeded the ratio expected from a random selection (gray stars in **Fig 3E**). This finding suggests that our pipeline successfully filters out priors not supported by the transcriptional data (20, 27).

### Combining sequence-based predictions of mRNA-sRNA interactions with transcriptomics data using the *Inferelator*

We showed above that the Inferelator is robust to the presence of false negatives and positives in the network priors (**Fig. 3**), so we exploited this property to separate true from false positives among sRNA-mRNA interactions that were predicted computationally. The strategy described below combines a sequence-based prediction step with a transcriptional data-driven filtering step (**Fig. 4A**). For any sRNA of interest, we first build a set of priors using a sRNA-mRNA interaction prediction method. Then, we run the Inferelator and recover the most likely targets of that sRNA. We chose CopraRNA (32) for the assembly of the sRNAs priors because it is a state of the art RNA-RNA interaction prediction method (9). It also offers an excellent framework to evaluate the potential of our method. A standard CopraRNA output contains 100 predictions (ranked by the associated p-values). CopraRNA performs a functional enrichment analysis among predicted targets. There is, however, no standard strategy to select which putative interactions should be investigated further. Any CopraRNA output will most likely include false positives that cannot easily be discarded. Therefore, our pipeline helps in selecting the most biologically relevant interactions among CopraRNA predictions.

**Figure 4.**
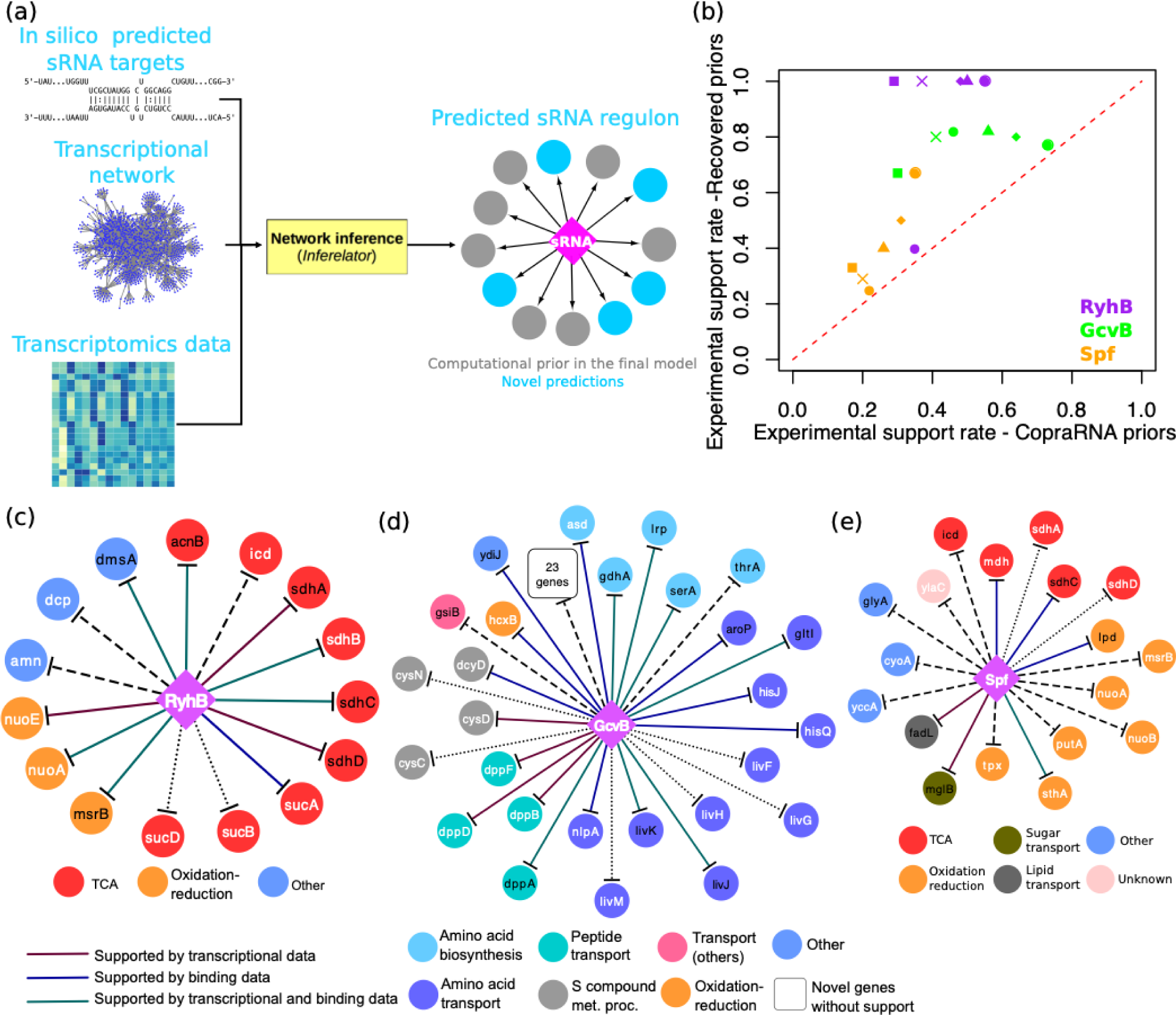
The Inferelator can identify computationally predicted sRNA-mRNA interactions with experimental support. A) General strategy to integrate computational sRNA-mRNA predictions in our pipeline. The resulting sRNA regulon is then analyzed to identify sequence-based sRNA-mRNA interactions supported by transcriptional data and potential additions to the sRNA regulon. B) The experimental support rate of recovered priors is significantly higher than the one of the original CopraRNA sRNA priors. The six points per sRNA correspond to the six sets of sRNA priors derived from CopraRNA predictions (**Table S2**). C) Inferred RyhB regulon when CopraRNA predictions associated with enriched functional terms were used as priors. D) Inferred GcvB regulon when CopraRNA predictions with p-value ≤ 0.01 were used as priors. E) Inferred Spf regulon when CopraRNA predictions with p-value ≤ 0.01 and associated with enriched functional terms were used as priors. Diamonds and circles represent sRNAs and target genes, respectively. Solid lines indicate interactions with experimental support. Dashed lines indicate interactions with no experimental support; dotted lines indicate unsupported targets that are part of an operon that contains experimentally supported targets. Priors included in the final regulon are labeled with black text. Novel targets (i.e. not present in the priors) are labeled with white text. Targets genes are colored based on their functional annotation.

We focused our analysis on the CopraRNA predictions for RyhB, GcvB and Spf. These sRNAs regulate different cellular processes and transcriptional profiling data indicate that each may directly or indirectly regulate dozens of genes. We hypothesized that if we used CopraRNA predictions as priors, our downstream activity estimation and network inference method would further distinguish between true and false positives and thus detect novel interactions. From the available transcriptional profiling data, we estimated that about 25% of the CopraRNA predictions are experimentally supported (i.e. differentially expressed when expression of the corresponding sRNA is perturbed or detection of physical interaction between sRNA and predicted targets; **Table S2**). To avoid a bias in our analyses, we compared five filtering strategies to reduce the proportion of unsupported priors in the initial set of CopraRNA predictions (**Table S2**). For each sRNA, we ran our pipeline using the following sets of priors: i) the full set of CopraRNA predictions. ii) targets with p-values ≤ 0.01. iii) targets associated with enriched functional terms. iv) the intersection of (ii) and (iii). v) the union of (ii) and (iii). vi) the union of the top 15 targets based on p-value (suggested in the original CopraRNA paper) and (iv). Experimental support rate of generated priors ranges from 0.17 to 0.73.

Initially, we compared the experimental support rates of the multiple sets of CopraRNA priors (generated with the above filtering strategies) to the inferred sRNA regulons. We observed that, in general, running the Inferelator dramatically shrank the initial set of priors (**Table S2**), while the experimental support rate increased significantly (**Fig. 4B**). This result supports the hypothesis that our method filters out false priors. Remarkably, we identified 26 sRNA-mRNAs predicted interactions that are most likely true additions to the corresponding *E. coli* sRNA regulons (**Table 2**). Each of the sRNA-mRNA interactions is supported by the transcriptional compendium analyzed with our network inference strategy, and independent experimental data (physical binding, transcriptional profiling or validation in a closely related species such as *Salmonella*). For example, the RyhB-*cheY* interaction is supported by the physical interaction between RyhB and *cheY* in *E. coli* and significant up-regulation of *cheY* in a *Salmonella* strain missing one of its two RyhB genes (10, 33). Another interesting target of *E. coli* RyhB is *mrp*. This interaction is supported by: 1) physical interaction between RyhB and *mrp* in *E. coli* (10); 2) increased translation rate of *mrp* in a RyhB deletion *E. coli* strain (31); and 3) the fact that *mrp* encodes an iron binding protein, which is consistent with the well-known role of RyhB in the iron sparing response (24). Therefore, the interactions listed in **Table 2** constitute a promising starting point for future experimental validation efforts.

**Table 2.**
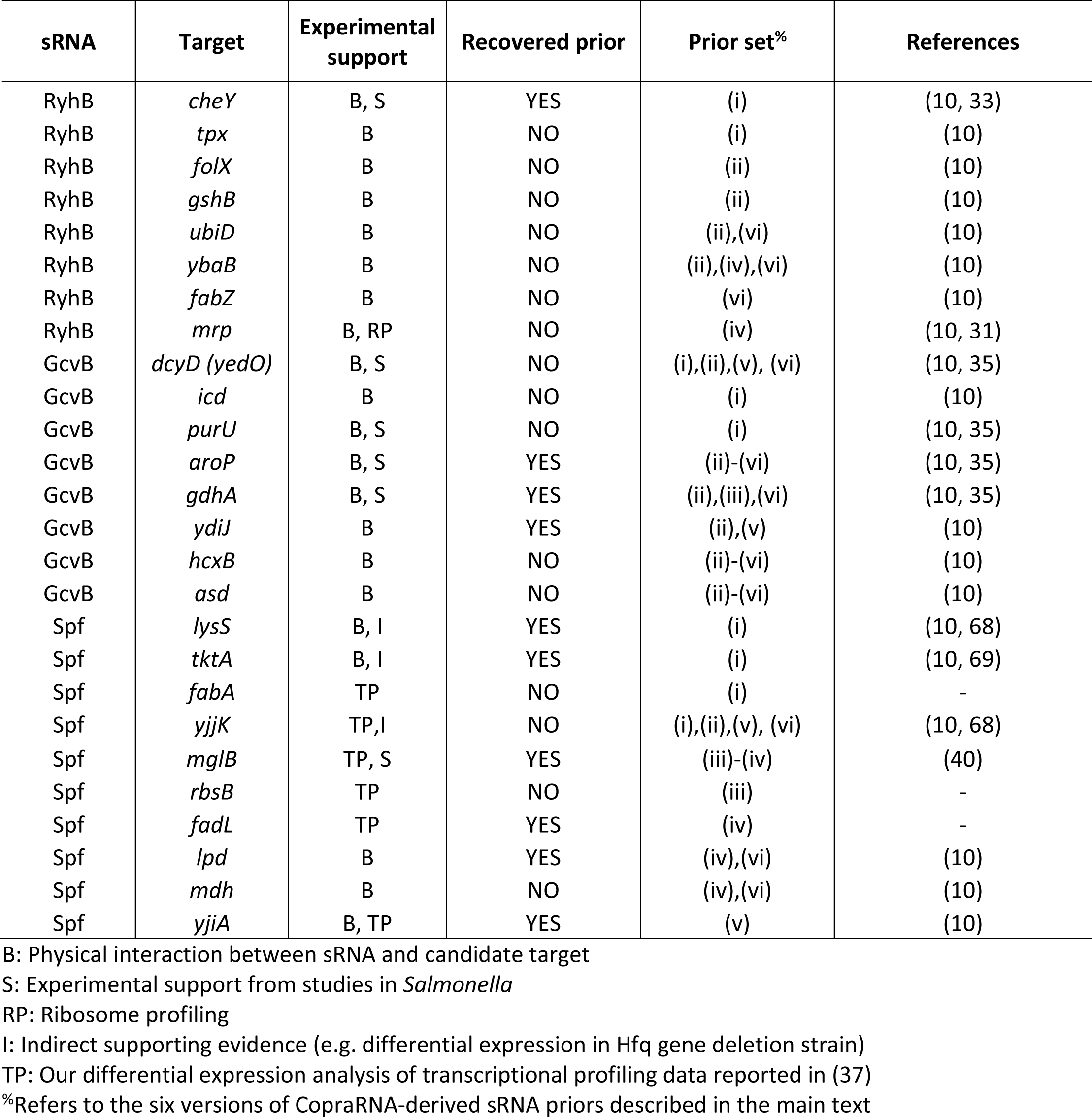
Putative new members of the RyhB, GcvB and Spf regulons identified using CopraRNA-derived sRNA priors.

Among the six sets of priors that we tested for RyhB, the one containing 38 genes associated with enriched functional terms gave the best results (**Fig. 4C**). Not only were all six priors included in the inferred network experimentally supported, but nine additional targets were predicted. Four out of the nine novel predictions had experimental support. Two additional targets (*sucB* and *sucD*) are in the same operon as *sucA*, one of the novel targets supported by binding data. Thus, the inferred RyhB regulon has a 0.67 accuracy (i.e. 10 out of 15 predicted targets are experimentally supported). The novel predictions (not present in the priors) included genes involved in respiration (*nuoA* and *nuoE*) and the citric acid (TCA) cycle (*sucA-sucB-sucD*), two cellular processes already associated with RyhB.

The five largest sRNA regulons inferred using CopraRNA-derived priors were for GcvB. Each of these GcvB regulons had 46 or more predicted targets (**Table S2**). This large size agrees with the global regulatory role of GcvB (34). **Fig. 4D** shows the inferred GcvB regulon when CopraRNA predictions with p-value ≤ 0.01 were used as priors. Eleven priors (out of 46) were recovered as putative targets in the inferred network. Nine out of those eleven priors had experimental support. In addition, 39 genes were predicted as novel targets. Although some of the novel predictions for GcvB lacked experimental support (23 out of 39), we identified multiple novel targets that show the potential of our approach. For instance, *nlpA* (validated as a GcvB target in 2018) (34) was predicted as a novel GcvB target. In fact, *nlpA* was predicted as a GcvB target in five of the six inferred GcvB regulons. Additionally, *asd*, *hisJ*, *hisQ*, *hcxB* and *dcyD* were novel predictions supported by physical binding data (10). The interaction between GcvB and *dcyD* was experimentally validated in *Salmonella* (35). *hisJ* and *hisQ* are in the same operon as *argT*, a known GcvB target included in the prior set. Four members of the *dpp* operon, involved in peptide transport (36) were predicted as additional GcvB targets; however, *dppA*, the first gene in the operon, was present among the priors. The inferred GcvB regulon included other six genes that belong to operons with known GcvB targets but lacked experimental support (dotted lines in **Fig. 4D**).

Predictions for Spf illustrate the performance of our method when a set of genes with low experimental support rate is used as priors. Spf had the lowest experimental support rate in each of the six versions of CopraRNA-derived priors (**Table S2**). **Fig 4E** shows the Spf regulon inferred using as priors the 23 genes predicted as Spf targets by CopraRNA with a p-value ≤0.01 and associated with enriched terms. Only eight of these 23 priors were experimentally supported. Yet, four out of the six recovered priors had experimental support. One of the novel targets was *sthA*, an experimentally validated target of Spf (37). An interesting prediction for Spf was *mdh*. The Spf-*mdh* interaction is supported by physical binding data and it is in line with the role of Spf in carbon metabolism (10, 37).

### Expanding the partially characterized sRNA regulons for GcvB, Spf, PrrF and FsrA

Here, we describe four examples to highlight how our pipeline successfully identified 15 novel sRNA-mRNA interactions with experimental support in *E. coli*, *P. aeruginosa* and *B. subtilis*. The predicted *E. coli* Spf regulon (from the inferred sRNA network in **Fig. 3**) is shown in **Fig. 5A**. Spf mainly controls genes associated with sugar metabolism and transport (37). Four (out of 12) priors were recovered as targets. Five additional targets were predicted, including *maeB*, which encodes a NADP-dependent malate dehydrogenase that converts malate into pyruvate (38). Originally, this gene was not reported as differentially expressed in a *spf* over-expressing strain (37). However, when we reanalyzed the transcriptional data with a Bayesian t-test, we found that *maeB* was significantly down-regulated in the over-expression strain. Moreover, a physical interaction between Spf and *maeB* mRNA was recently reported by Melamed et al (10). We conclude that *maeB* is a true novel Spf target. The NAD-dependent malate dehydrogenase of *E. coli*, *maeA*, is a known Spf target (37, 38). Interaction of Spf with *maeB* and *maeA* (located in independent transcriptional units) indicates that Spf can completely block the generation of pyruvate from malate by repressing both types of malate dehydrogenases. Another novel predicted target for Spf was the *mgl* operon. *mglB* encodes a galactose ABC transporter (39) and it was significantly down-regulated in the *spf* over-expressing strain (exclusively detected in our analysis). *mglB* is predicted as a Spf target by CopraRNA (**Fig. 4E**) and the Spf-*mglB* interaction was recently validated in *S. enterica* (40). Considering that *E. coli* and *S. enterica* are phylogenetically close, we conclude that the *mgl* operon is also a true Spf target in *E. coli*. This implies that Spf represses galactose metabolism (through the repression of members of the *gal* operon) (41) and transport of galactose into the cell (by repressing the *mgl* operon). The fifth novel Spf target was *sdhA*. This prediction is supported by the validated interaction between Spf and *sdhC*, the first gene of the *sdh* operon (42). Desnoyers and Massé found that Spf primarily regulates the *sdh* operon at the translational level. Thus, the inclusion of the Spf-*sdh* interaction in our model indicates either that our approach can detect interactions producing subtle changes in mRNA stability (indeed, Desnoyers et al. observed degradation of the *sdh* mRNA 30 mins after Spf induction) or that Spf induces a faster degradation of the *sdh* polycistronic mRNA in a still unidentified condition. Overall, we found that all five of the Spf novel targets were experimentally supported.

**Figure 5.**
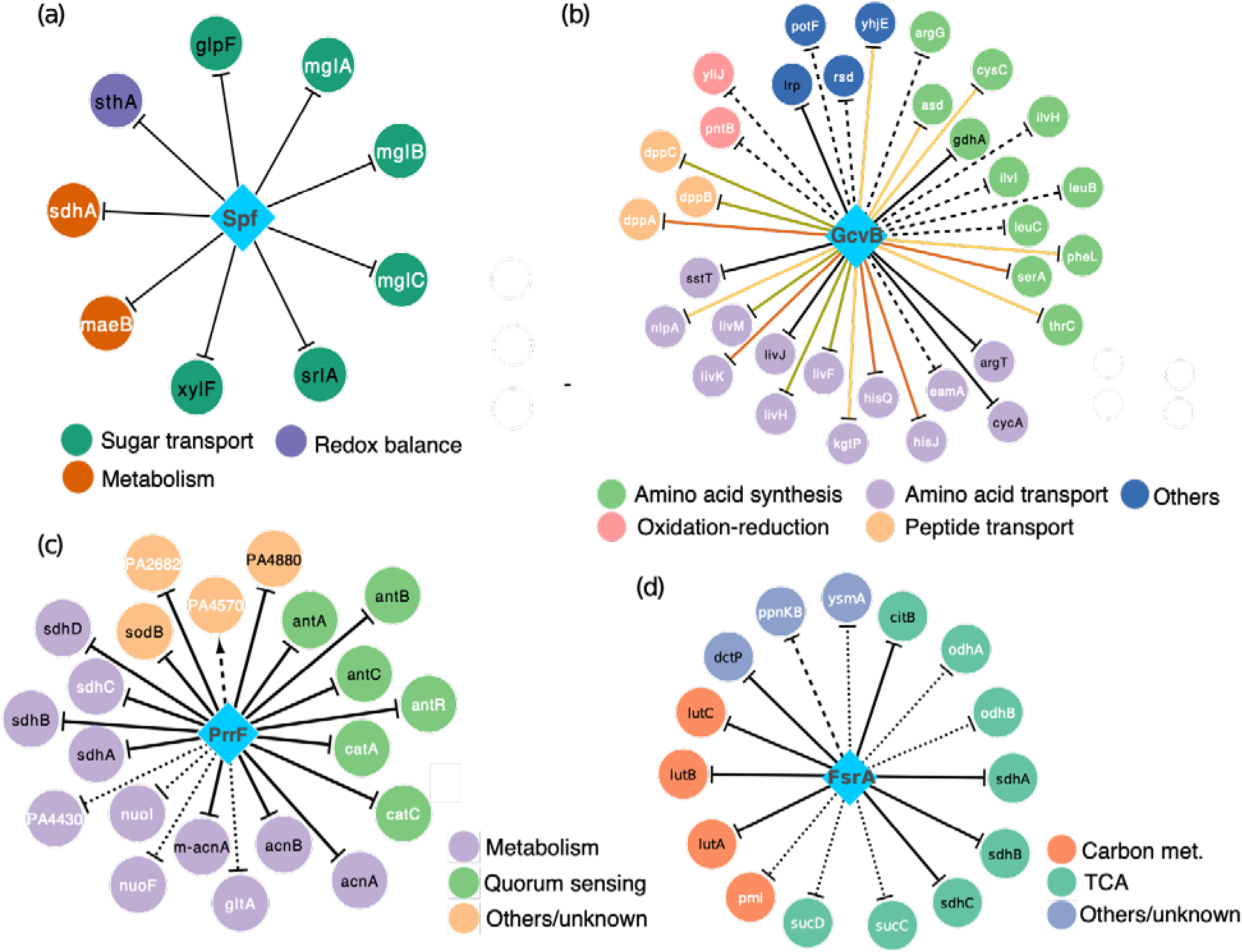
Selected expanded sRNA regulons of *E. coli*, *P. aeruginosa* and *B. subtilis*. White node labels indicate novel targets (not present in the sRNA priors) and black node labels indicate prior targets. Solid lines indicate priors and experimentally supported novel targets. Dotted lines indicate interactions partially supported by experimental data, computational RNA-RNA prediction methods or data from functional analogs in other species. Dashed lines indicate unsupported predictions. A) The inferred *E. coli* Spf regulon. All predicted targets are experimentally supported. B) The inferred *E. coli* GcvB regulon. Novel interactions supported by transcriptional profiling data, physical binding data or both are shown in green, orange and red, respectively. C) The inferred PrrF regulon of *P. aeruginosa*. D) The inferred FsrA regulon of *B. subtilis*. In each panel, nodes are color-coded by functional categories.

The inferred *E. coli* GcvB regulon, using the compiled set of experimentally supported sRNA priors, is shown in **Fig. 5B**. Our model expands the role of GcvB in the regulation of amino acid biosynthesis and transport. Six (out of nine) priors were recovered as GcvB targets and 27 additional targets were predicted. Ten of the novel targets (not included in the prior sRNA network) were supported by transcriptional profiling data. Among these novel targets, several have been validated, i.e. *serA* [validated in *S. enterica* (34, 35)]; three members of the *dpp* operon (*dppA*, *dppB*, *dppC*) (43); four members of the *livKHMGF* operon (*livF*, *livH*, *livM* and *livK*), and two members of the *argT-hisJQMP* operon (*hisJ* and *hisQ*). The set of novel GcvB targets has significant experimental support from physical binding data (10) (12 out 27 new targets, for a hyper-geometric test p-value= 2.7 × 10^−11^). Notably, there were five novel targets (*asd*, *kgtP*, *nlpA*, *pheL*, *yhjE*) not supported by transcriptomics data, but detected *in vivo* by Margalit and collaborators as physically interacting with GcvB (10). Two predicted GcvB targets (*thrC* and *cysC*) were indirectly supported by binding data (10). GcvB interacts with *thrL* (10), the leader peptide sequence of the *thr* operon that contains *thrC*. GcvB physically interacts with *cysB* (10), encoding the transcriptional regulator of *cysC*. To the best of our knowledge, assigning to the GcvB regulon *asd* (also predicted as a target when CopraRNA-derived priors were used), and *kgtP*, which respectively encode an aspartate-semialdehyde dehydrogenase and an α-ketoglutarate H^+^ symporter (44, 45), is only supported by our model and physical binding data collected in (10). One of the main challenges of new technologies capturing physical binding between sRNAs and mRNAs [e.g. RNA Interaction by ligation and Sequencing (10), *in vivo* UV crosslinking with RNA deep sequencing (40), MS2-Affinity Purification coupled with RNA Sequencing (34)] is to identify whether those interactions do actually influence mRNA stability and translation rate (10). Thus, our approach constitutes a complementary tool to identify which interactions, among the hundreds of detected binding events, have functional relevance.

PrrF1 and PrrF2 are two iron-responsive sRNAs of *P. aeruginosa* and function as analogs of *E. coli* RyhB (46). Both sRNAs are transcriptionally repressed by Fur under iron rich conditions (46). PrrF1 and PrrF2 are almost identical at the sequence level and are adjacent on the *P. aeruginosa* chromosome. Thus, they were considered as a single regulator (PrrF) in our analysis. The predicted PrrF regulon is shown in **Fig. 5C**. All eleven priors were recalled in the final model and the inferred PrrF regulon included 10 novel targets. Five genes (*antR*, *catA*, *catC*, PA2682 and *sdhC*) were significantly up-regulated in a wild-type *P. aeruginosa* strain grown in iron rich medium (compared to the WT grown under iron poor conditions, i.e. when PrrF is active) and in the double *prrF1*-*prrF2* deletion mutant (compared to the WT strain) (46). We considered those five targets to be experimentally supported. Independent regulation of *antR* and its target *antABC* operon was validated in (47). *sdhC* can also be considered a validated target per (46). *gltA*, which encodes an enzyme involved in the TCA cycle, was significantly up-regulated in the WT strain grown in iron rich medium vs. the iron poor condition (47). Since PrrFs are repressed at high iron concentrations, the observed up-regulation of *gltA* supports our prediction. Moreover, *gltA* is a known target of FsrA (a functional analog of PrrF) in *B. subtilis* (48). For the reasons described above and the involvement of known PrrF targets in the TCA cycle, we considered *gltA* to be a probable PrrF target. We hypothesize that the novel targets *nuoF*, *nuoI* and PA4430 are likely regulated by PrrF despite not being supported by available transcriptional profiling data. We base our conclusion on the following observations: first, interaction of PrrF with the *nuo* operon and the PA4427-PA4428-PA4429-PA4430-PA4431 operon is supported by the PrrF1-*nuoK* and PrrF1-PA4431 interactions predicted by CopraRNA. Second, the predicted PrrF*-nuo* interaction is highly probable considering that: i) the *nuo* operon is regulated by RyhB in *E. coli*; ii) PrrF and RyhB have multiple targets in common (e.g. the *sdh* operon, *acnA* and *acnB*). The same is true for the PrrF-PA4430 interaction. PA4430 putatively encodes cytochrome b and RyhB regulates multiple cytochrome encoding genes. Third, we considered as differentially expressed the genes labeled as such in (47). In that study, a 0.0001 p-value threshold was used to define differential expression. The stringent p-value threshold may account for categorizing PA4430 and the *nuo* operon as not differentially expressed. Hence, only one out of the 10 novel PrrF targets predicted by the model (PA4570) appears to be a false positive. Other putative targets are supported either by experimental data, computational RNA-RNA predictions, or conservation of PrrF targets in other species.

FsrA is the functional analog of PrrF in *B. subtilis* (25). **Fig. 5D** shows the inferred FsrA regulon. The predicted regulon contains eight (out of twelve) priors and seven novel target genes. In agreement with our model, three novel targets (*odhA*, *odhB* and *pmi*) are among the FsrA targets predicted by CopraRNA. *odhA* and *odhB* form an operon and encode genes involved in the TCA cycle (49). In support of the predicted FsrA-*odhA* interaction, analysis of 2D protein gels showed that average fold-change of the OdhA protein level is 1.78 in the double *fsrA*-*fur* deletion mutant respect to the WT strain (25). Furthermore, *odhA* and *odhB* mRNA levels were up-regulated (1.68 and 1.91 mean fold-changes, respectively) in the double *fsrA*-*fur* deletion mutant when compared to the *fur* single deletion mutant (48). The interaction between FsrA and *ysmA*, an uncharacterized gene, is supported by the similarity between *ysmA* transcription profile and that of known FsrA targets (*leuC* and *sdh* operon) (48). Additionally, *ysmA* is up-regulated (3.78 mean fold-change) in the double *fsrA*-*fur* deletion mutant with respect to the *fur* deletion mutant (48). The interaction between FsrA and the *sucC*-*sucD* operon is supported by up-regulation (2.46 mean fold-change) of *sucD* in the double *fsrA*-*fur* deletion mutant compared to the *fur* single deletion mutant (48) and down-regulation (0.53 mean fold-change) of *sucC* in the *fur* single deletion mutant compared to the WT (48). Two predicted targets of FsrA (*sucC* and *ppnKB*) are validated targets of RoxS, another trans-encoded sRNA of *B. subtilis* (17). Interestingly, expression of multiple genes regulated by Fur, the transcriptional repressor of FsrA, also appear to be influenced by RoxS (17). This may suggest a functional connection between RoxS and FsrA. However, we did not find any data in support of the predicted FsrA-*ppnKB* interaction. Follow up experiments are required to obtain a definitive answer. In summary, interactions between FsrA and the *odhA*-*odhB* and *sucC*-*sucD* transcripts are particularly promising due to the role played by these genes in the TCA cycle, a previously known target of FsrA (*sdh* operon) (25, 48).

## CONCLUSIONS

We have developed a new computational pipeline that integrates estimates of TF and sRNA activities with a well-tested network model selection procedure for inferring bacterial sRNA regulons. Our work shows that using transcriptional profiles of sRNAs as proxy for their activity in traditional network inference approaches is less than optimal, because it does not account for the fact that sRNA activity can be influenced by factors such as RNA chaperones, ribonucleases and sRNA: targets ratio (18, 19). Our findings further demonstrates that the need to estimate regulatory activity of non-coding RNAs is not exclusive to eukaryotic systems (50) but relevant for all types of regulatory non-coding RNAs that require substantial processing and are involved in multiple interactions (such as micro-RNAs and bacterial sRNAs).

Our results indicate that integration of sRNA activities in network inference pipelines significantly improves their prediction power (**Fig. 3A**) and our strategy significantly outperforms previous network inference efforts. Importantly, this work complements sRNA-mRNA prediction methods based on sequencing analysis and the recently developed technologies for detecting physical interactions between sRNAs and mRNAs (**Fig. 4**). Our computational approach identified a total of 39 novel sRNA-mRNA interactions with experimental support in Gram-positive and Gram-negative species (*E. coli*, *P. aeruginosa* and *B. subtilis*). In addition, we showed that our strategy is robust to false positives and negatives, thus allowing the accurate detection of novel sRNA targets. Importantly, our method is especially well suited to removing the many false positives present in sequence-based computationally predicted sRNA-mRNA interactions. Our pipeline can both expand current sRNA regulons and serve as a first approach to prioritize the study of predicted targets of uncharacterized sRNAs.

The sRNA regulons inferred in this study increase by 40% the number of experimentally supported interactions originally compiled for estimating *E.* coli sRNA activities. We uncovered novel experimentally supported sRNA-mRNA interactions (**Fig. 4**, **Fig. 5A-B** and **Table S2**) involved in chemotaxis and oxidation-reduction pathways. Thus, our work extends the contribution of sRNA-mediated regulation in these processes. Simultaneously, we discovered how a single sRNA (Spf) can repress all branches of a metabolic reaction (i.e. conversion of malate to pyruvate in NAD and NADP dependent fashion). Analysis of the inferred Spf regulon also suggested how sRNAs can repress the consumption of alternative sugars (i.e. galactose) by simultaneously inhibiting their catabolism and their intake. In general, our approach offered insights into the functional role of bacterial sRNAs as fine tuners of gene expression in the analyzed species.

The main limiting factor in our approach is the fact that it requires prior information (including transcriptomics data, a transcriptional network, and candidate sRNA targets). As a proof of principle, we selected bacterial species for which we could comprehensively assess the quality of the inferred models. Beyond these selected species, we believe that there is a much larger group of bacterial species (e.g. *Salmonella enterica* and *Mycobacterium tuberculosis*), whose study could benefit from the application of the strategy delineated in this work. Transcriptional compendia for approximately 20 bacterial species can be easily downloaded from the COLOMBOS database (51). Transcriptional networks can be (at least partially) reconstructed by mining literature and databases that store information about experimentally supported transcriptional interactions [e.g. RegPrecise database (30)]. Finally, we show that initial sets of sRNA priors can be generated using available mRNA-sRNA interaction prediction tools (e.g. CopraRNA), genetic perturbations or with global detection of sRNA-mRNA binding events.

The applicability of our strategy will increase in the next few years as the field of bacterial sRNA-mediated regulation grows. Incorporating estimated TFs regulatory activity in network inference strategies, has led to recent improvements in the transcriptional regulatory networks of yeast (52), sex specific gene networks in *Drosophila* (53), transcriptional networks associated with cancer (54, 55) and transcriptional networks that drive differentiation of mice T lymphocytes (56). Our strategy relies on knowledge about sRNA-mRNA interactions that is already available to accurately estimate sRNA activities and to identify novel sRNA targets. Hence, we expect performance of our strategy to improve as the quality and number of confirmed sRNA-mRNA interactions continues to rise.

## MATERIALS AND METHODS

### Bacterial species

We inferred transcriptional regulatory networks and small non-coding RNA regulons for *Escherichia coli*, *Pseudomonas aeruginosa, Staphylococcus aureus* and *Bacillus subtilis*.

### Small non-coding RNA priors

sRNA-mRNA interactions used as sRNAs priors for sRNA activity estimation in each species are listed in **Table S1**. For sRNA priors in *E. coli*, only one member of each operon containing multiple validated sRNA targets was considered to avoid over-representation of any operon. Because S596 is an uncharacterized sRNA of *S. aureus*, we used as S596 priors the CopraRNA-derived candidate targets selected in (26).

### Transcriptomics datasets

The transcriptomics datasets used for inferring the transcriptional and sRNA networks of analyzed species are described in **Table S3**.

### Prior transcriptional networks

For each species, the prior transcriptional network was constructed as a collection of experimentally supported signed (activation or repression) TF-gene interactions. The prior networks were used for estimating the regulatory activities of TFs included as potential regulators, inferring the corresponding transcriptional network and defining the final model of the inferred networks (*see below*). Sources for each species are shown in **Table S3**.

### Estimation of transcription factors and sRNAs regulatory activities

Activities of potential regulators (TFs and sRNAs) were simultaneously estimated using the set of experimentally supported interactions in the prior network as described in (27). Briefly, we first combined the sRNA and transcriptional prior networks into a global prior network. We represented the analyzed transcriptional dataset in matrix format (referred to as ***X***) where each row corresponded to the transcriptional profile of a gene. Then, we applied a network component analysis (NCA; 43) to decompose ***X*** in two matrices: a first matrix ***P***, which we derived from the prior network. The values in ***P*** are in the {0, 1, −1} set, where 1 and −1 indicate activation and repression, respectively. Value in the *P_ij_* entry corresponds to the interaction between gene *i* and regulator *j*. The second matrix ***A*** is unknown but represents the activities of regulators along the conditions in ***X***. As such, the *A_kl_* entry is the activity of regulator *k* in condition *l*. In matrix notation, NCA can be stated as: ***X****=**PA*** (Eq. 1). We solved for ***A*** using the pseudo-inverse of ***P*** as explained in (27).

### Inference of Transcriptional and sRNA Networks

Transcriptional and sRNA networks were simultaneously inferred using Inferelator *B*ayesian *B*est *S*ubset *R*egression (BBSR), as detailed in (27). The core model of the Inferelator with incorporation of TFs and sRNAs activities can be summarized as: *x_i,j_* = ∑*_k ∈{TFs U sRNAs}_ β_i,k_*Â*_k,j_* (Eq. 2), where *x_i,j_* is the mRNA level of gene *i* in condition *j*, Â is the matrix of estimated activities generated with NCA (as described above), and *β_i,k_* indicates the effect (positive or negative) and strength of regulator *k*’s activity on gene *i*. *β* is the main output of the Inferelator. To model the sparsity of biological networks, BBSR solves for a matrix *β* where most values are zero. More details about BBSR solution can be found in (27). To avoid overfitting, we bootstrapped the input transcriptional data 20 times (we have previously observed minimal change above 20 bootstraps) (27). We averaged the *β* scores associated with each re-sampling instance into a final *β* matrix. The second output of the Inferelator, is a confidence score matrix generated as explained in (27). The confidence score of an interaction indicates the likelihood of the interaction. Mixed-CLR was run using the *mi_and_clr.R* script in the Inferelator release, available in https://sites.google.com/a/nyu.edu/inferelator/home.

### Construction of final model of transcriptional and sRNA networks

We ranked the set of all potential regulator (TF/sRNA)-gene interactions based on the associated confidence scores. We used a 0.5 precision cutoff [as previously used in (27)] to determine the set of interactions included in the final model. The confidence cutoff was defined as the score at which exactly 50% of the TF-gene and sRNA-gene interactions above the cutoff were part of the prior network.

### Validation of inferred sRNAs regulons

For each species, we mined publicly available transcriptional profiling data, sRNA-mRNA binding data and results of other relevant experiments (such as northern blots, point mutations, translational fusions, ribosome profiling, *in-silico* predictions) for assessing the accuracy of the inferred sRNA regulons. A total of 385 candidate *E. coli* sRNA-mRNA interactions were suggested by available literature (excluding binding data). This set of potential interactions was extended to 691 with the addition of genes located in the same operons. *E. coli* operons prediction was downloaded from MicrobesOnline (58). Independent studies supporting novel sRNA-mRNA interactions discussed in the text are cited in the relevant sections.

### Differential expression analysis of Spf over-expressing E. coli

Normalized microarray data of Spf over-expression (GEO accession GSE24875) (37) was downloaded and differential expression analysis was performed using a Bayesian T-test with Cyber-T (59). Only genes included in the *E. coli* transcriptomics data used in this study were considered in the analysis. In addition, genes that were absent in any of the replicates were excluded. Finally, genes with p-values ≤ 0.01 were considered differentially expressed. We have successfully used this p-value threshold for analyzing *B. subtilis* data (27).

### In-silico prediction of sRNA-mRNA interactions

For sRNAs that were conserved among multiple bacterial species, precomputed predictions from the CopraRNA website (http://rna.informatik.uni-freiburg.de/CopraRNA/Input.jsp) were downloaded and used as priors. If CopraRNA predictions were not available for a sRNA of interest, a new run was submitted to the CopraRNA website. All CopraRNA predictions were downloaded between January and June 2016.

### Functional enrichment analysis

Enrichment analysis was performed on the DAVID website (60).

## ACKNOWLEDGEMENTS

We thank Wade Winkler for critical discussions about this project and comments of the manuscript. We are grateful to Jonathan Goodson for critical reading of the manuscript. This work was supported by the Simons Foundation (RB), the National Institute of Health (5T32AI100853; R01-DK103358-01 and R01-GM112192-01 to RB) and by funds from the Zegar Family Foundation (to PE). The funders had no role in study design, data collection and interpretation, or the decision to submit the work for publication.

## AUTHORS CONTRIBUTIONS

MLAO, RB and PE designed research. MLAO, CH and BS performed research. MLAO, NSB, RB and PE analyzed data. MLAO, RB and PE wrote the paper.

---

**Figure S1.**
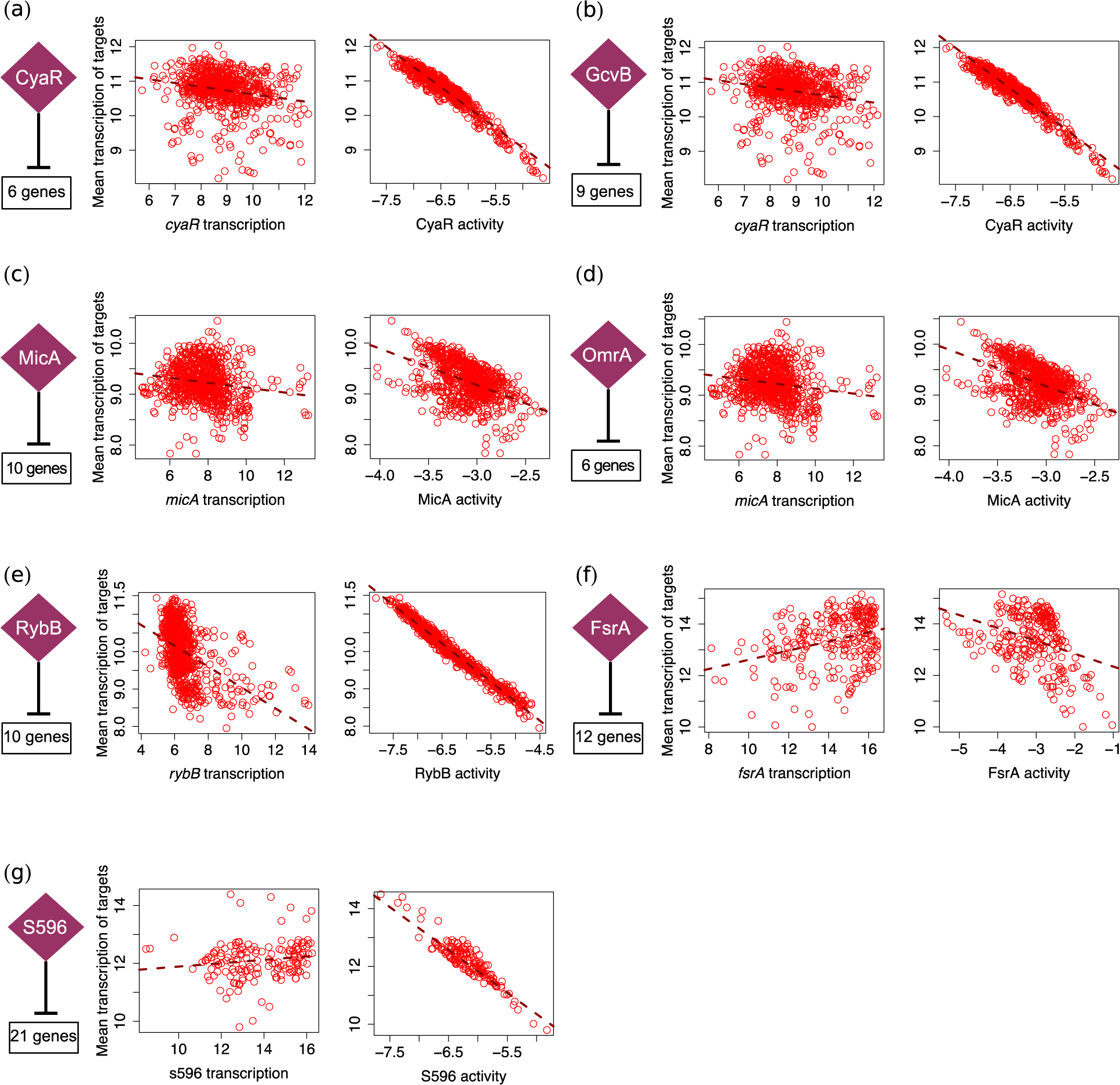
Motivation for estimating the regulatory activity of sRNAs in Gram-positive and Gram-negative bacteria. sRNA activities were estimated for each experimental condition. Each dot represents one microarray experiment. The number of experimentally supported targets used to estimate sRNA activities and to compute the mean transcription of the analyzed regulons (in each condition) is indicated. In all cases, a stronger relation is observed between the estimated sRNA activities and the average transcription profile of their dependent genes (right panels) than between the transcription profile of the sRNAs and the average transcription profile of their targets (left panels). A) *E. coli* CyaR controls genes involved in sugar metabolism (63). CyaR is expressed during high cellular levels of cAMP. B) *E. coli* GcvB regulates genes involved in amino acid transport and amino acid biosynthesis (64). C) *E. coli* MicA is a stress related sRNA (65). D) *E. coli* OmrA is important in the response to membrane stress (66, 67). E) *E. coli* RybB is a stress related sRNA (65). MicA and RybB have multiple targets in common. F) FsrA is involved in the iron sparing response of *B. subtilis* (25). FsrA is a functional analog of RyhB in *E. coli*. G) S596 was recently identified as the functional analog of RyhB and FsrA in *S. aureus* (26).

**Figure S2.**
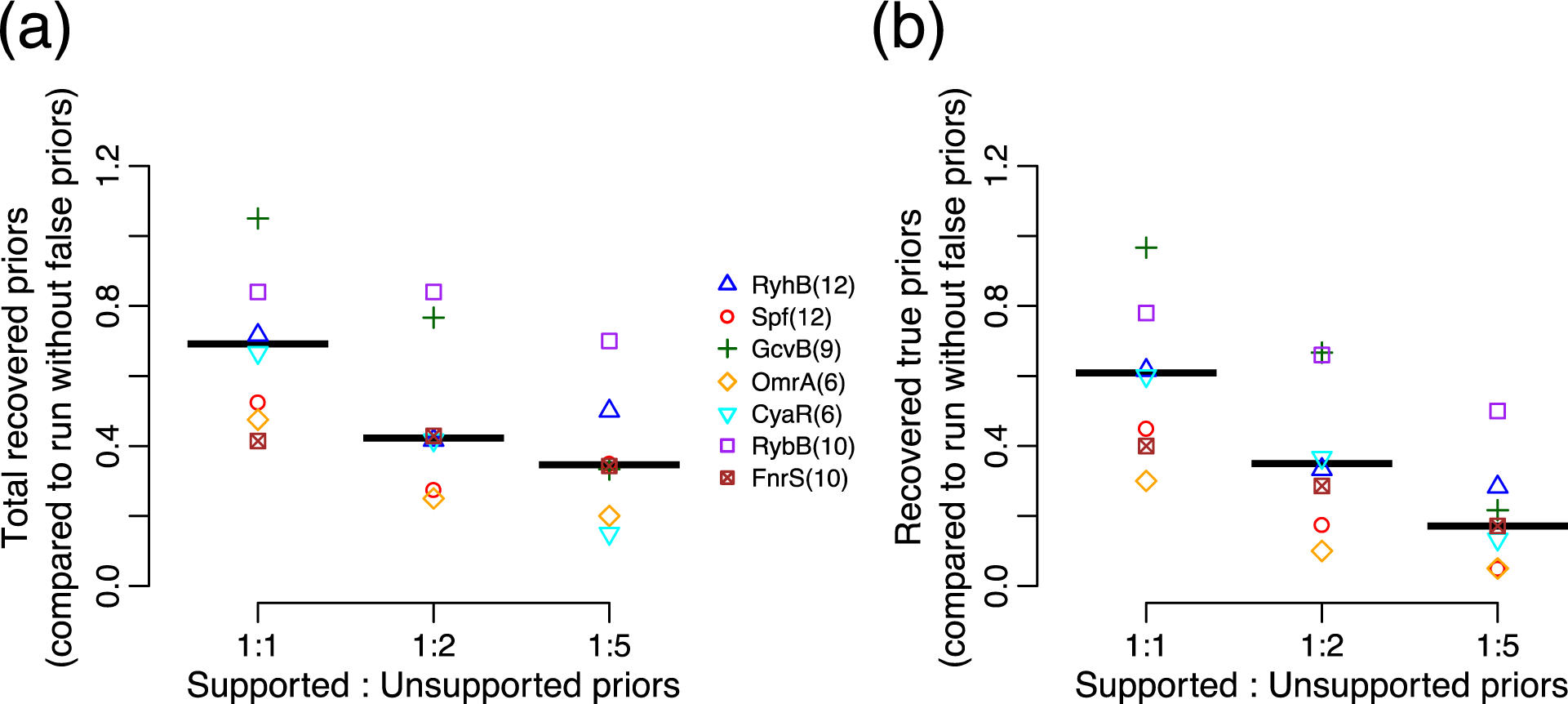
Presence of false sRNA-mRNA interactions reduced the number of sRNA priors included in the networks inferred by the Inferelator. Each dot is the mean value of ten Inferelator runs (each one with a different set of false priors). Each colored symbol corresponds to one of seven sRNAs. Black lines indicate the median proportion for all seven sRNAs. Number of true targets for each sRNA is shown in parentheses. A) Ratio between the number recovered priors in Inferelator runs with noisy sRNA priors and the total number of recovered priors in the Inferelator run without false positives. B) Ratio between number of recovered priors with experimental support (true priors) in Inferelator runs with noisy sRNA priors and the total number of recovered priors in the Inferelator run without false positives.

**Table S1.**
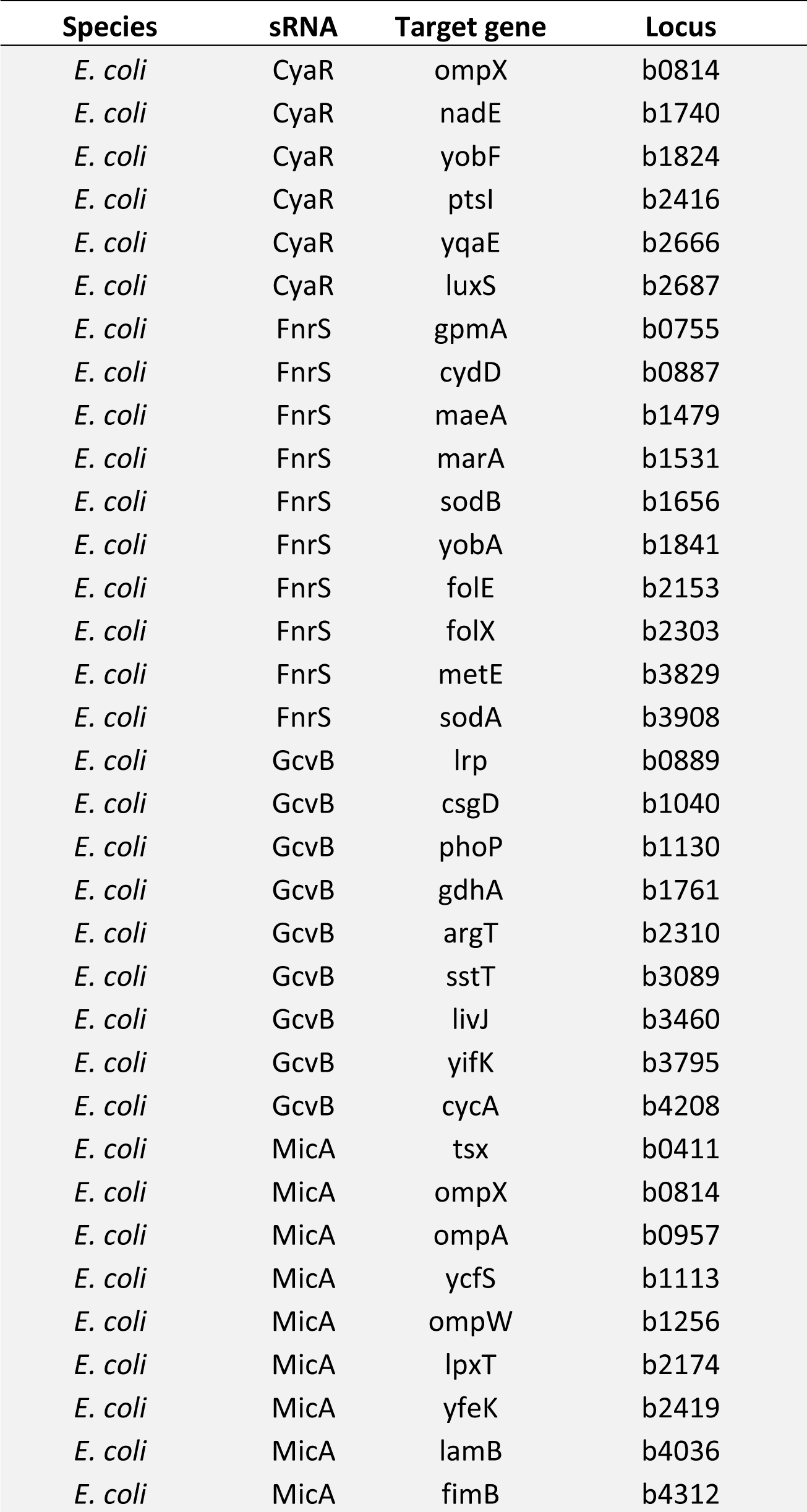
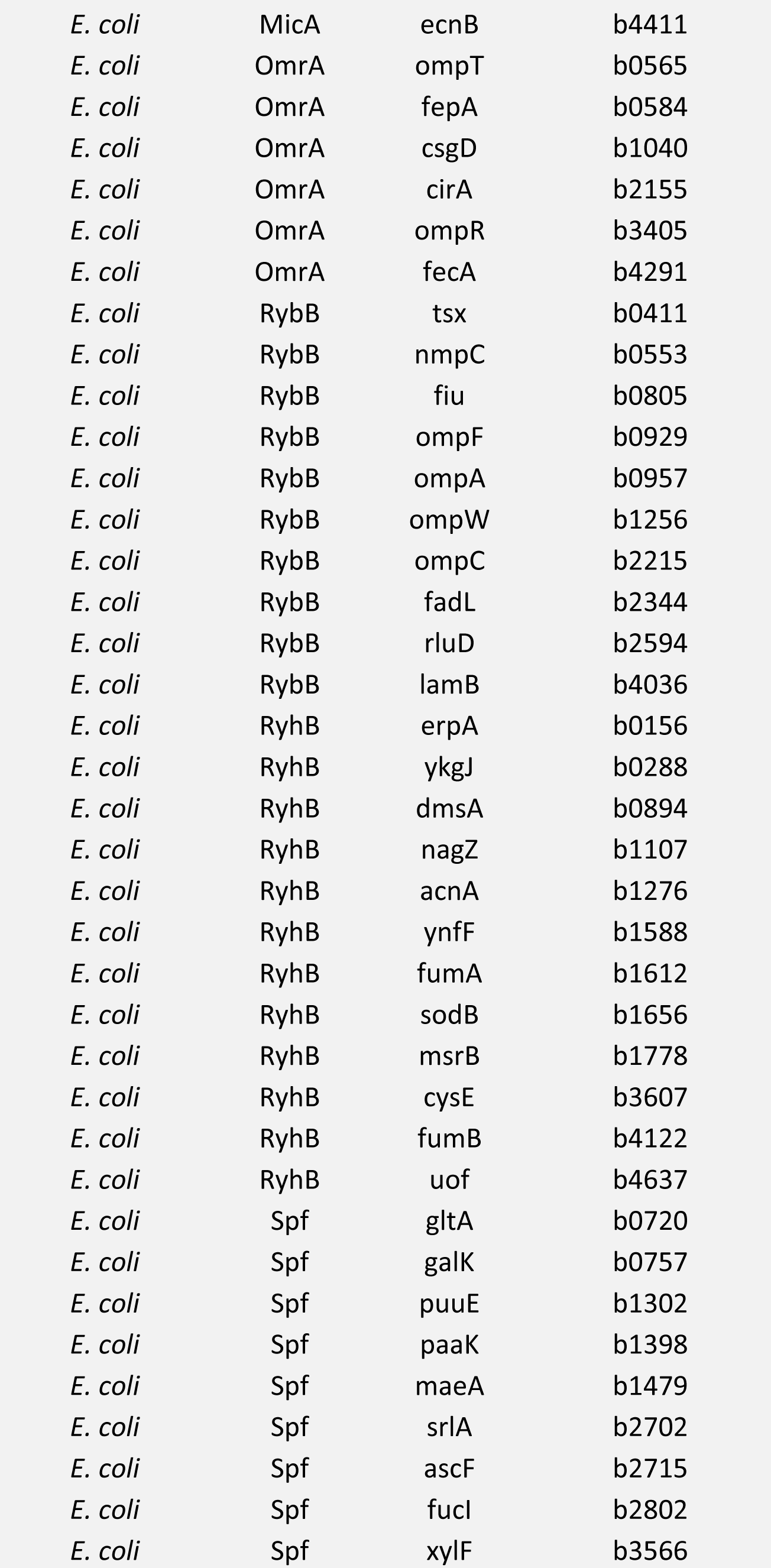
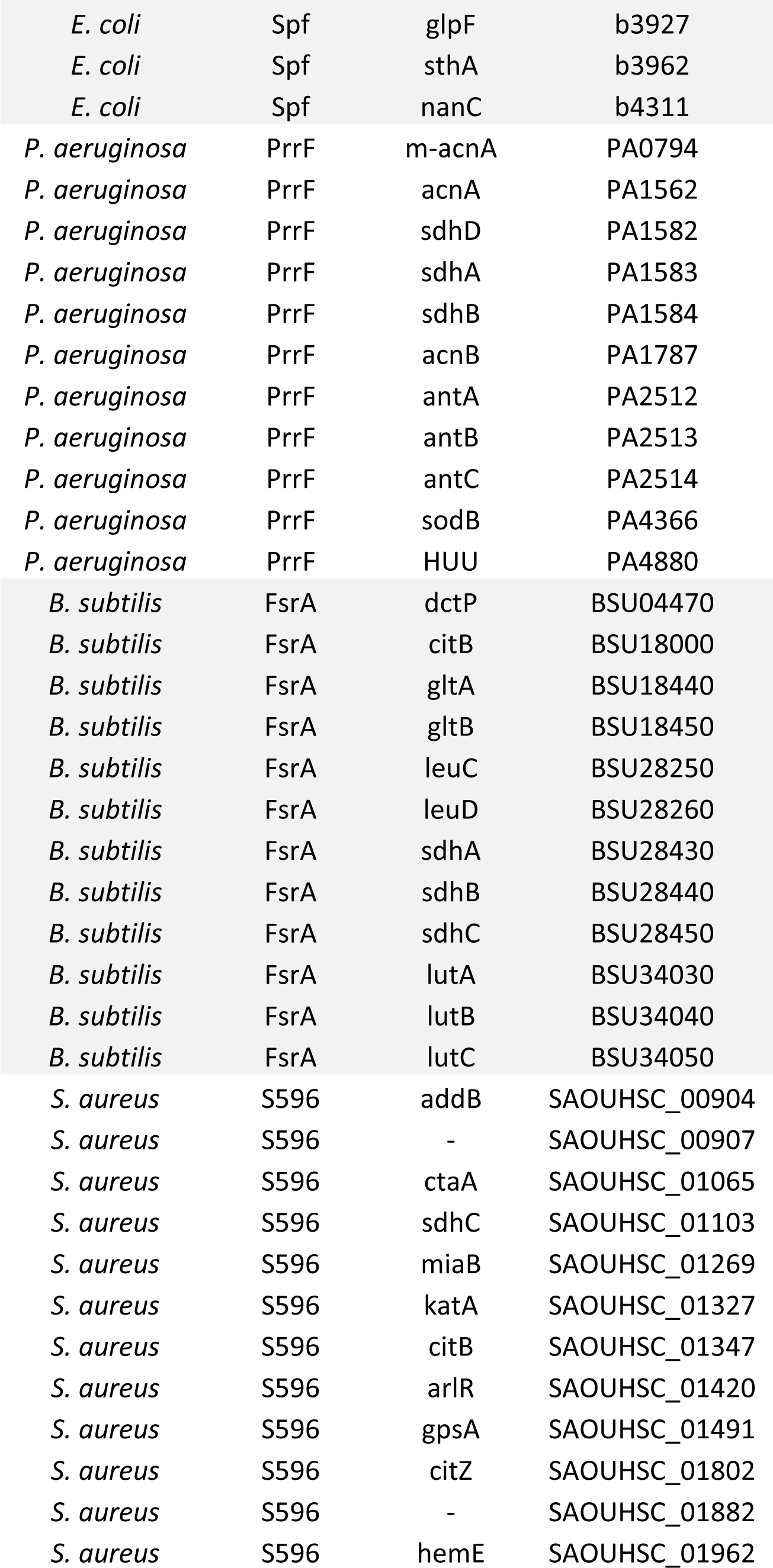
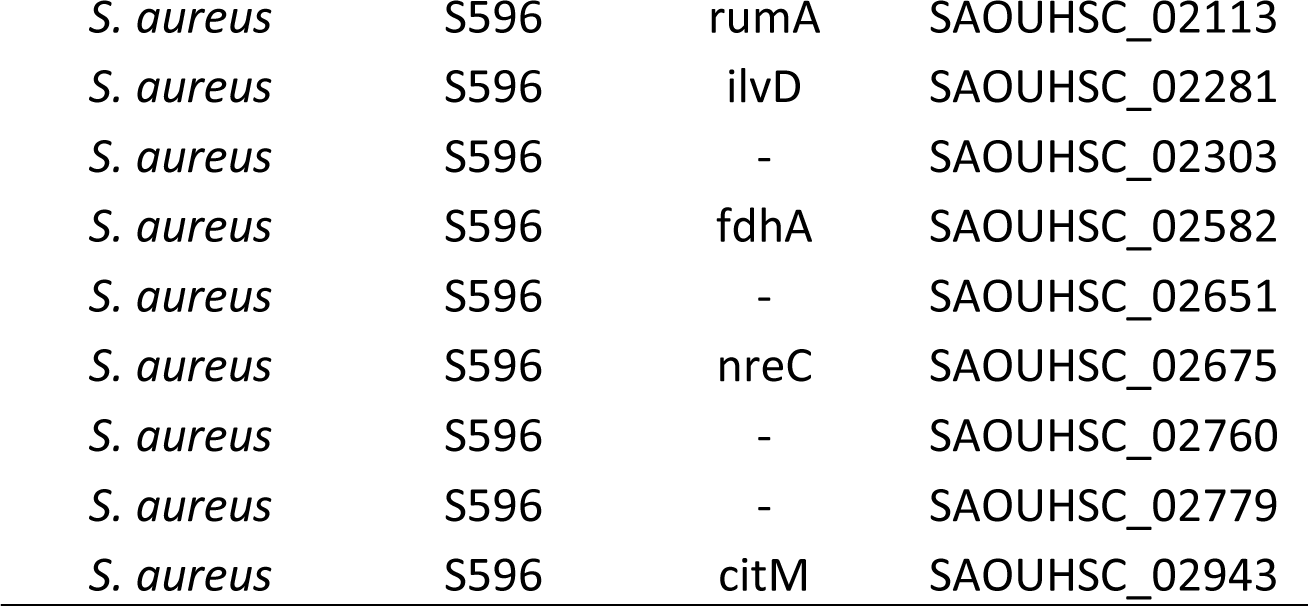
sRNA-mRNA interactions used as priors in this study.

**Table S2.** The Inferelator filters the CopraRNA-derived priors and predicts novel sRNA-mRNA interactions with experimental support.

| sRNA Prior selection<sup>&</sup> | sRNA | Priors | Supported priors<sup>#</sup> | Priors predicted as targets<sup>#</sup> | New targets<sup>#</sup> |
| --- | --- | --- | --- | --- | --- |
| Top 100 (i) | RyhB | 100 | 29 (0.29) | 5 (1) | 6 (0.17) |
|  | GcvB | 100 | 30 (0.3) | 3 (0.67) | 12 (0.42) |
|  | Spf | 100 | 17 (0.17) | 6 (0.33) | 27 (0.07) |
| P-values $\leq 0.01$ (ii) | RyhB | 49 | 17 (0.35) | 5 (0.4) | 29 (0.14) |
|  | GcvB | 46 | 21 (0.46) | 11 (0.82) | 39 (0.41) |
|  | Spf | 54 | 12 (0.22) | 4 (0.25) | 17 (0.06) |
| Annotated with enriched terms (iii) | RyhB | 38 | 19 (0.5) | 6 (1) | 9 (0.44) |
|  | GcvB | 34 | 19 (0.56) | 11 (0.82) | 43 (0.37) |
|  | Spf | 43 | 11 (0.26) | 5 (0.4) | 6 (0.17) |
| P-values $\leq 0.01$ AND annotated with enriched terms (iv) | RyhB | 20 | 11 (0.55) | 1 (1) | 6 (0.33) |
|  | GcvB | 22 | 16 (0.73) | 13 (0.77) | 37 (0.46) |
|  | Spf | 23 | 8 (0.35) | 6 (0.67) | 12 (0.17) |
| P-values $\leq 0.01$ OR annotated with enriched terms (v) | RyhB | 67 | 25 (0.37) | 4 (1) | 1 (0) |
|  | GcvB | 58 | 24 (0.41) | 10 (0.8) | 39 (0.36) |
|  | Spf | 74 | 15 (0.2) | 7 (0.29) | 17 (0.06) |
| Top 15+<sup>§</sup> (vi) | RyhB | 29 | 14 (0.48) | 1 (1) | 8 (0.38) |
|  | GcvB | 25 | 16 (0.64) | 10 (0.8) | 36 (0.5) |
|  | Spf | 32 | 10 (0.31) | 4 (0.5) | 14 (0.14) |

Predicted targets with physical interaction with sRNAs according to binding data in (10), or differential expression in transcriptional profiling experiments (overexpressing or deleting putative sRNA regulator) were considered supported. Members of operons with differential expression in sRNA perturbation were also considered experimentally supported.
<sup>&</sup>Derived from CopraRNA predictions
<sup>#</sup>Experimental support rate is shown in parentheses
<sup>§</sup>Prior set was the union of the top 15 predictions (ranked by associated p-values), and the set of targets with p-values $\leq 0.01$ and annotated with enriched terms (iv)

**Table S3.** Transcriptional prior networks and transcriptomic datasets used in this study

| Species | Transcriptional prior network<sup>%</sup> | Reference | Transcriptomic dataset<sup>#</sup> | Reference |
| --- | --- | --- | --- | --- |
| *Escherichia coli* | RegulonDB version 9.0 (1875<sup>*</sup>) | (29) | Many Microbe Microarrays<sup>†</sup> (4297 x 861) | (13) |
| *Pseudomonas aeruginosa* | Collection of known transcriptional interactions | (70) | COLOMBOS 3.0<sup>Δ</sup> (5629 x 559) | (51) |
|  | Experimental sigma factor network | (71) |  |  |
|  | RegPrecise<sup>&</sup> (3569<sup>^</sup>) | (30) |  |  |
| *Staphylococcus aureus* | SigB regulon | (26) | HG001 (2837 x 156) | (26) |
|  | RegPrecise<sup>&</sup> (798<sup>^</sup>) | (30) |  |  |
| *Bacillus subtilis* | SubtiWiki | (72) | BSB1 (4445 x 269) | (73) |
|  | Transcriptional network constructed with the Inferelator (2614<sup>Σ</sup>) | (27) |  |  |

<sup>%</sup>Number of interactions in the corresponding network is shown in parentheses <sup>#</sup>Number of genes and number of arrays in the dataset are shown in parentheses <sup>\*</sup>Only signed interaction with strong or confirmed evidence were included as priors <sup>†</sup>Version with un-averaged replicates was used. 16 conditions related to sRNA KOs or their regulators were removed <sup>&</sup>Only signed interactions (activation or repression) were considered <sup>Δ</sup>Total number of interactions in the compiled network (including all mentioned sources) <sup>Δ</sup>123 rows missing more than 10% of their values were removed <sup>Σ</sup>Total number of interaction the compiled network. The network is composed of TF-gene interactions originally reported in SubtiWiki (72) that were recovered in the network reconstructed by the Inferelator, and experimentally supported novel interactions of the Inferelator-reconstructed model (27)

